# Environmental statistics and sensory experience shape patch foraging strategies in *Drosophila* larvae

**DOI:** 10.64898/2026.03.27.714746

**Authors:** Akhila Mudunuri, Klára Tučková, Ahmed El Hady, Katrin Vogt

## Abstract

Animals foraging in patchy environments must balance exploiting current resources with exploring for better alternatives to maximize resource intake and to survive. However, the neural and computational mechanisms underlying such adaptive decisions have just recently begun to be understood. Using *Drosophila* larvae as an experimentally tractable model, we combine long-timescale behavioral tracking in controlled patchy environments with varying statistics, along with quantitative analysis and computational modeling, to dissect foraging decision strategies. We show that larvae flexibly adjust their behavior according to both the quality and valence of available resources, shaped by prior foraging experience. A simple integration model recapitulates larval patch-leaving behavior, with model parameters tuned by environmental statistics and foraging history. Together, these findings establish *Drosophila* larvae as a powerful system for studying adaptive foraging and for uncovering the neural circuit mechanisms that implement experience-dependent foraging decisions.

## Introduction

Foraging is essential for the survival of all organisms, as food is necessary for growth, maintenance, and reproduction. However, in natural environments, food quality varies dynamically across space and time, making it challenging to adjust foraging decision strategies. Therefore, animals must continuously balance the energetic costs of exploration against the nutritional gains of exploitation to decide when to leave a resource patch (1). Classical foraging models, such as the marginal value theorem (2), predict that an animal should leave a resource patch when the instantaneous intake rate falls below the average intake rate of the environment. However, this relies on the unrealistic assumption that animals have access to global environmental information and that the environmental statistics are stable over time (3, 4).

Effective foraging requires animals to infer local resource quality and flexibly adapt their foraging behavior in response to environmental changes (5). Empirical studies have shown that prior experience influences foraging decisions in diverse taxa, including crabs (6), bees (7), ants (8), and mammals (9, 10). To account for experience-dependent effects, an updated foraging model was proposed in which recent foraging history modulates the timing of departure from the current patch (11). However, this model assumes an imposed, fixed, and finite memory timescale for updating the reward rate; it does not explicitly treat patch-leaving as a decision-making process and excludes potential uncertainties in the foraging environment. Alternatively, one can address these limitations using a drift diffusion modeling framework to model patch-leaving decisions as an evidence-accumulation process in which patch departures are driven by noisy integration of evidence (12). Patch departure is triggered when a decision variable crosses a threshold. Importantly, recent experimental studies have shown that cortical neural activity reflects integration dynamics associated with patch-leaving behaviors (13, 14).

With a powerful genetic toolkit and an available whole-brain connectome, *Drosophila* larvae are an excellent model for dissecting the neural mechanisms underlying foraging decisions and for directly relating behavioral function to neural structures. Larvae feed continuously to reach the critical weight required for pupation (15). During development, larvae must acquire carbohydrates for energy (16) and proteins for growth and development (17). The balance between these macronutrients strongly influences developmental rate and adult lifespan (18). In nature, however, larvae grow on decaying plant or fungal matter that is scarce and distributed in discrete, patchy resources (19), necessitating efficient inter-patch foraging strategies. Previous work has shown that larvae exhibit higher inter-patch foraging when there is no protein in their current patch (20) and can compensate for nutrient deprivation during development by changing their feeding behaviors (21). Larvae adapt their locomotor behavior in response to various patch substrates (22, 23). However, the foraging study by Wosniack et al. was conducted in spatially homogeneous environments using odor-rich substrates with very different nutritional profiles, making it difficult to disentangle the effects of resource quality, nutritional value, and olfactory navigation.

Here, we address this limitation by using a well-controlled behavioral assay that tracks larval foraging over hours-long timescales in patchy environments containing resources with similar nutritional profiles and no odor cues. This controlled approach allows us to reliably dissect the foraging decision strategies of fly larvae across a variety of environmental structures. We found that larval foraging behaviors depend on both resource quality and valence. Larvae flexibly adjust their foraging strategies based on the resources they encounter, and these strategies are shaped by prior foraging experience. Together, our results demonstrate that larval foraging is context and history-dependent. Moreover, computational modeling indicates that larval foraging behavior is mediated by an integration mechanism that optimizes patch residence time depending on the quality or value of the patch and previous patch experience.

## Results

### Larval foraging behavior depends on resource quality

To understand how larval foraging behavior is affected by resource quality, larvae were tested in a large square arena (25 cm x 25 cm) containing 100 ml of 2% agar. Four circular patches (radius: 1.75 cm) were embedded in the arena at equal distances (6.25 cm) from the center (Fig. 1A). Each patch consisted of 1.5 ml of 2% agarose mixed with either 0.1M (Movie S1) or 1M fructose (Movie S2), concentrations that are attractive to larvae (24–26). Patches were designed to be large enough to function as a non-depleting resource, while still requiring larvae to explore the arena to locate the patches. In this “homogenous foraging environment”, all patches contain the same resource. Pure agarose patches were used in control experiments to assess potential patch boundary effects (Movie S3). Individual early third-instar larvae were placed in the center of the arena. Their behavior was recorded continuously for 3 hours and analysed using custom tracking software.

**Figure 1.**
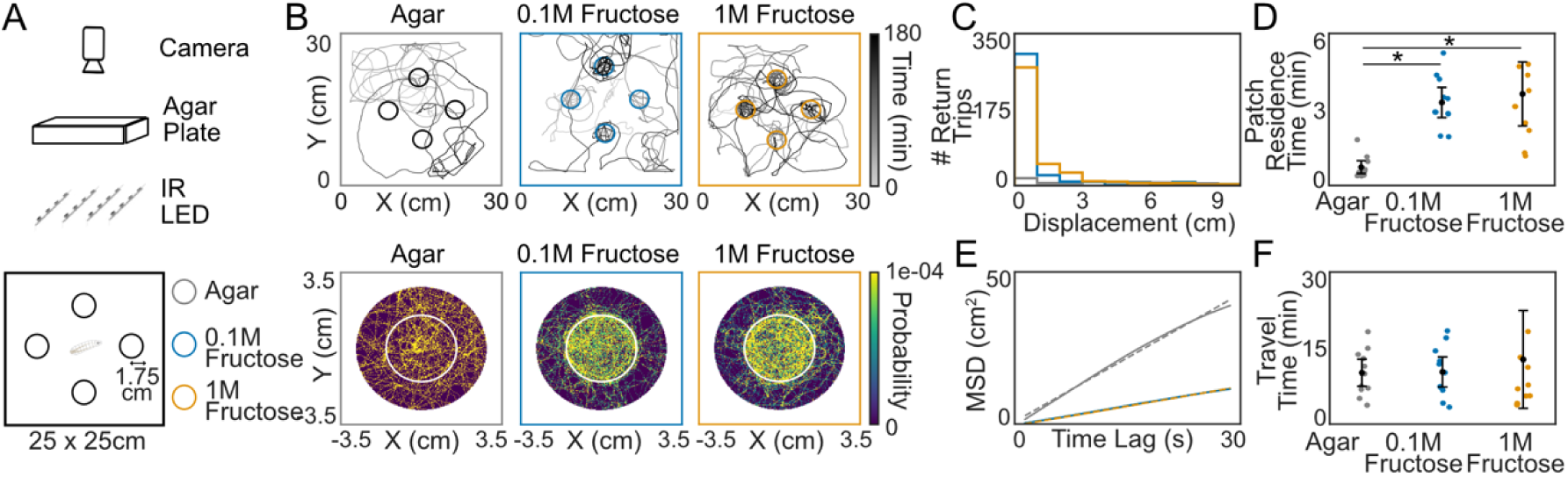
*Drosophila* larval foraging depends on resource quality: (**A**) Individual larvae were placed in the middle of a square arena (25 x 25 cm) containing 2% agar and 4 patches of the same resource (agar (gray), 0.1M fructose (blue), 1M fructose (yellow)). The behavior of a single larva was recorded for 3 hours under infrared illumination. (**B**) Sample trajectories of individual larvae for 3 hours when the patches contained agar, 0.1M fructose, or 1M fructose. The patch heatmap visualizes the spatial distribution of larvae both on the patch (marked by a white circle) and in the surrounding area across all patches (N = 11 larvae per patch condition). (**C**) Number of return trips to the patch as a function of the maximum displacement from the patch edge after the exit. (**D**) Average thresholded patch residence time for each larva. The dots represent individual larvae, and the line shows the mean ± 95% confidence interval. (Mann-Whitney U test, * p < 0.05) (**E**) Mean squared displacement (MSD) of larvae on-patch as a function of time lag. The dotted line indicates the linear fit to the MSD (Agar: 1.3636, 0.1M Fructose: 0.41401, 1M Fructose: 0.40739). (**F**) Average inter-patch travel time of the larvae.

Most larvae visited all patches (Fig. S1A) and exploited patches containing fructose (Fig. 1B). They also spent time in the close vicinity of patches containing fructose, likely due to resource diffusion (27) or larvae overshooting the patch boundaries. One of the most important behavioral measures in patch foraging is patch residence time, defined as the duration a larva spends on a resource patch. We calculated the average patch residence time for each larva. Larvae had a longer patch residence time on fructose patches (Fig. S1B). There was no difference in patch residence time on 0.1M fructose patches compared to 1M fructose patches.

As larvae often remain near fructose patches after exiting and seem to return after a short time, we quantified both their displacement from the patch edge and the travel time required to return. We observed a correlation between maximum displacement and time spent outside the patch (Fig. S1C), suggesting that displacement can serve as a threshold for including short trips in the patch residence time (Fig. S1D). A displacement threshold of 3 cm was chosen, which included the majority of return trips made by larvae (Fig. 1C). Before applying the threshold, larvae entered fructose patches more frequently than agar patches (Fig. S1E), as there was a higher number of return trips made by larvae to patches containing fructose. However, after applying the return trip threshold, the number of patch entries was similar across conditions. After thresholding, larvae showed even higher patch residence times on the fructose patches (Fig. 1D), with no difference between 0.1M and 1M fructose concentrations. The average patch residence time remained unchanged over the 3-hour experimental period in both fructose conditions, while it decreased in the agar condition (Fig. S1F).

Movement is an integral part of patch foraging as both spatial and temporal decisions are crucial to optimizing foraging strategies. To quantify larval movement dynamics on and off the patch, we calculated mean squared displacement (MSD) and run speed. The larvae had a lower MSD (Fig. 1E) and were slower on fructose patches (Fig. S1G). There was no difference in larval speed off-patch; however, larvae generally had a lower MSD on arenas containing fructose patches (Fig. S1H). Travel time, defined as the time required for larvae to move from one patch to another, was similar across conditions (Fig. 1F) and remained stable over time (Fig. S1I). We further quantified the time larvae spent in different parts of the arena. Larvae spent less time at the arena border when patches contained 1M fructose and more time in regions near the patches compared to 0.1M fructose and agar (Fig. S1J). The time spent in the center of the arena did not differ between the conditions.

### Larval foraging behavior depends on resource valence

As environments rarely contain only attractive resources, we investigated how resource patches with different valences influence larval foraging behaviors. Each patch consisted of 1.5 ml of 2% agarose containing either 0.1M (Movie S4) or 1M salt (Fig. 2A, Movie S5). Lower salt concentrations are attractive to larvae, whereas higher concentrations are aversive (28).

**Figure 2.**
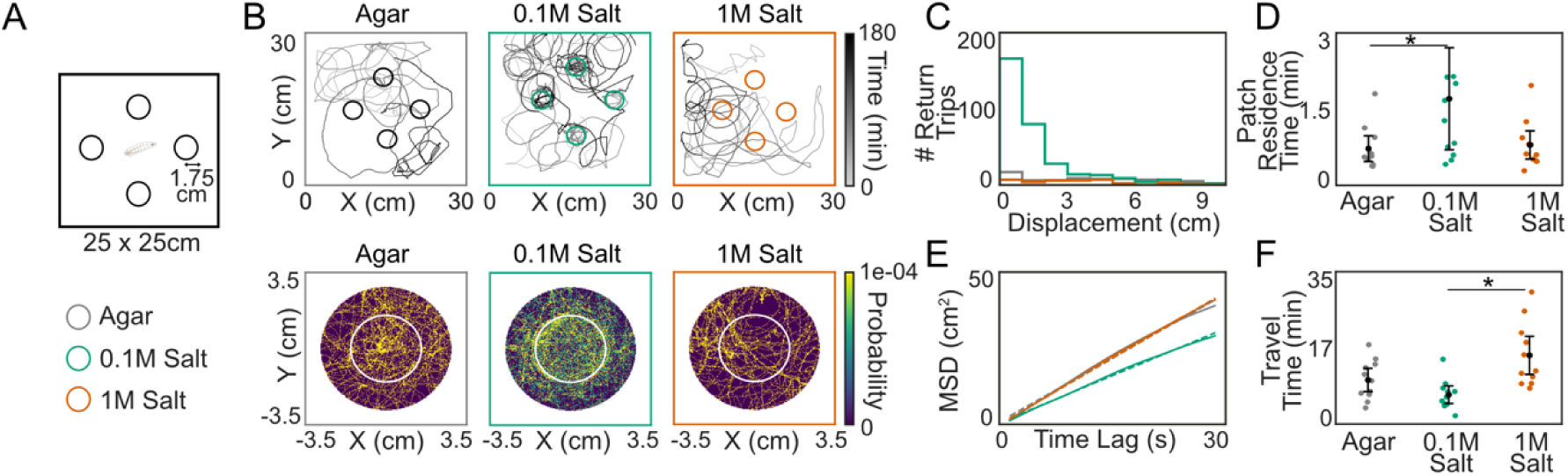
*Drosophila* larval foraging depends on resource valence: (**A**) Individual larvae were placed in the middle of a square arena (25 x 25 cm) containing 2% agar and 4 patches of the same resource (agar (gray), 0.1M salt (green), 1M salt (orange)). (**B**) Sample trajectories of individual larvae for 3 hours when the patches contained agar, 0.1M salt, or 1M salt. The patch heatmap visualizes the spatial distribution of larvae both on the patch (marked by a white circle) and in the surrounding area across all patches (N = 11 larvae per patch condition). (**C**) Number of return trips as a function of the maximum displacement from the patch edge. (**D**) Average thresholded patch residence time for each larva. Dots represent individuals, and the line shows the mean ± 95% confidence interval. (**E**) MSD of larvae on-patch; dotted line is the linear fit to the MSD (Agar: 1.3636, 0.1M Salt: 1.0026, 1M Salt: 1.4092). (**F**) Average inter-patch travel time of the larvae. (Mann-Whitney U test, * p < 0.05)

Most larvae visited all patches in the homogeneous foraging environments (Fig. S2A). Larvae exploited 0.1M salt patches more than 1M salt and agar patches (Fig. 2B). They also spent time in the vicinity of patches containing 0.1M salt, likely due to resource diffusion creating locally attractive salt concentrations or larvae overshooting the patch boundaries. However, there was no difference in the average patch residence time across conditions (Fig. S2C). Because larvae often stay in the vicinity of the patches after leaving, we quantified both their displacement from the patch edge and the travel time required to return to the patch. We found that the maximum displacement was correlated with the time spent outside the patch (Fig. S2C), indicating that displacement could be used as a threshold to classify brief trips as part of a patch visit (Fig. S2D). We used a displacement threshold of 3 cm, as it accounts for the majority of return trips to the patch (Fig. 1C).

Before applying the threshold, larvae entered the 0.1M salt patches more than the 1M salt and agar patches (Fig. S2E), as larvae made more return trips to patches containing 0.1M salt (Fig. 2C). However, after applying the threshold, the number of patch entries was lower for 1M salt than for 0.1M salt. Applying this threshold, we found that larvae spent more time on 0.1M salt patches during an average patch visit (Fig. 2D). The average patch residence time within the salt conditions did not differ during the 3-hour experiment (Fig. S2F).

The larvae had a lower MSD (Fig. 2E) and were slower on 0.1M salt patches compared to agar and 1M salt (Fig. S2G). However, off-patch larval speed and MSD (Fig. S2H) did not differ between the conditions. Travel time between patches was higher when the patches contained 1M salt compared to 0.1M salt and agar (Fig 2F). This may be because larvae spent more time in the vicinity of the 1 M salt patches, but did not enter them. Travel time did not change over time within each condition (Fig. S2I). The proportion of time spent at the arena border, near patches, and in the central region was comparable across the conditions (Fig. S2J).

### Larvae adapt their behavior based on resource quality and valence

Environments typically contain multiple resources, requiring organisms to adjust their behavior flexibly to maximize their food intake. To test whether larvae adapt their foraging behavior to varying resource qualities, we placed them in a “heterogeneous environment” containing food patches of two different resource qualities for 3 hours. Specifically, two patches contained low-quality resources (0.1M fructose), and two patches contained high-quality resources (1M fructose) (Fig. 3A, Movie S6).

**Figure 3.**
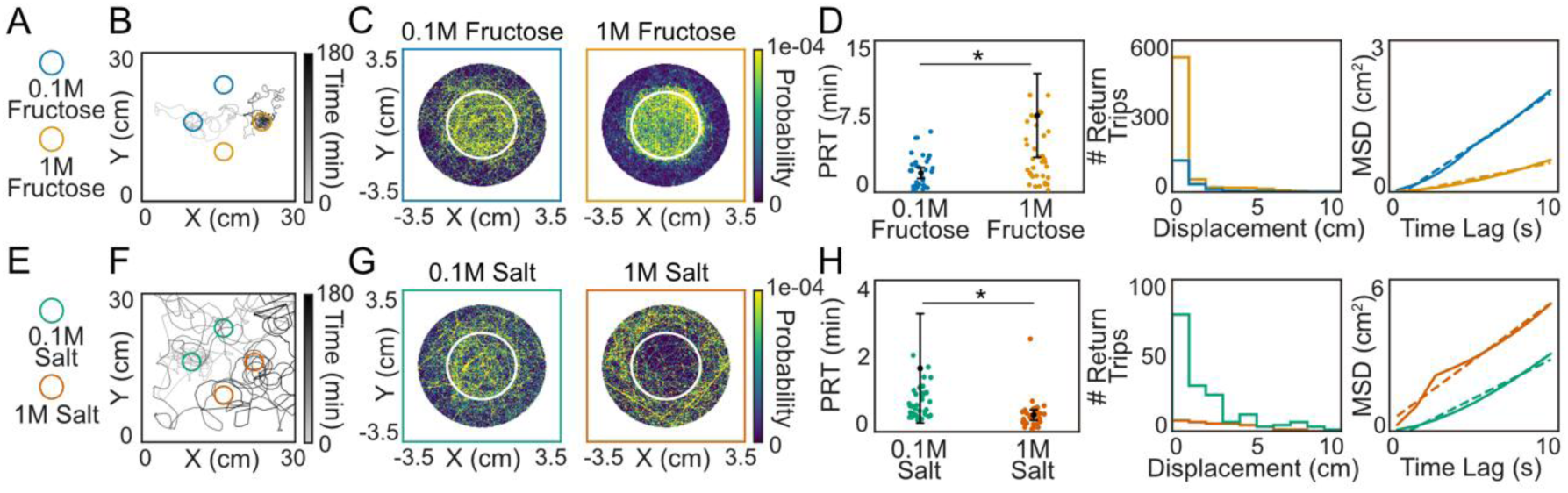
*Drosophila* larvae adapt their foraging behavior based on resource quality and valence: (**A**) Square arena (25 × 25 cm) containing 2% agar, with two 0.1M fructose patches (blue) and two 1M fructose patches (yellow). (**B**) Sample trajectory of an individual larva over 3 hours. (**C**) Patch heatmap showing the spatial distribution of larvae both on the patches (outlined in white) and in the surrounding arena for 0.1M and 1M fructose patches (N = 21 larvae). (**D**) Average thresholded patch residence time for each larva on 0.1M and 1M fructose patches. Dots indicate individual larvae, and the line represents the mean ± 95% confidence interval. Number of return trips as a function of the maximum displacement from the patch edge (Mann-Whitney U test, * p < 0.05). MSD of larvae on-patch. The dotted line indicates the linear fit to the MSD (0.1M Fructose: 0.256, 1M Fructose: 0.079). (**E**) Square arena (25 × 25 cm) containing 2% agar, with two 0.1M salt patches (green) and two 1M salt patches (orange). (**F**) Sample trajectory of an individual larva over 3 hours. (**G**) Patch heatmap for 0.1M and 1M salt patches (N = 21 larvae). (**H**) Average thresholded patch residence time for each larva on 0.1M and 1M salt patches. Number of return trips made by the larvae. MSD of larvae on-patch. The dotted line indicates the linear fit to the MSD (0.1M Salt: 0.383, 1M Salt: 0.559).

All larvae visited at least one low-quality and one high-quality fructose patch (Fig. 3B, Fig. S3A). Larvae spent more time on and in the vicinity of 1M fructose patches than on 0.1M fructose patches (Fig. 3C). To quantify this, we calculated the patch residence time (thresholded, thus including return trips) and found that larvae stayed longer on and returned more to high-quality fructose patches than to low-quality patches (Fig. 3D, Fig. S3B). Consistently, larvae made more thresholded entries onto 1M fructose patches than onto 0.1M patches (Fig. S3C). Over time, there was a decrease in patch residence time on 0.1M fructose patches; however, the patch residence time on 1M fructose patches did not change (Fig. S3D). Larvae exhibited lower MSD and slightly reduced speed on 1M fructose patches compared to 0.1M patches (Fig. S3E).

To examine whether larvae adjust their foraging strategies in response to resource valence, we exposed them to a heterogeneous environment arena containing salt patches of opposing valences for 3 hours. The environment consisted of two attractive low-salt patches (0.1M) and two aversive high-salt patches (1M) (Fig. 3E, Movie S7). All larvae sampled both patch types, visiting at least one attractive and one aversive salt patch during the experiment (Fig. 3F, Fig. S3F). Larvae showed a preference for 0.1M salt patches, spending more time on and near them while avoiding 1M salt patches (Fig. 3G). This preference was consistent with an increase in the thresholded residence time and number of return trips to 0.1M salt patches compared to 1M salt patches (Fig. 3H, Fig. S3G). Similarly, larvae had more thresholded patch entries onto 0.1M salt compared to 1M salt (Fig. S3H). Patch residence time increased over time for 0.1M salt patches, whereas it remained unchanged for 1M salt patches (Fig. S3I). Larvae displayed a lower MSD on 0.1M salt patches than on 1M salt patches, whereas the speed did not differ between the conditions (Fig. S3J).

### Larvae adapt their foraging behavior based on their past foraging experience

As larvae adapt their foraging behavior in response to available resources, we asked whether these changes arise from an immediate response to the current resource patch or prior foraging experience. To address this, we compared the patch residence times on the first and second patches encountered by the larvae in a heterogeneous patch foraging arena containing different fructose qualities (Fig. 4A). Similar to the homogeneous environment (Fig. 1), we observed a slight reduction in patch residence time when transitioning between patches of the same resource quality in the heterogeneous environment (Fig. 4B). In contrast, patch residence time increased when larvae transitioned from a low-quality patch (0.1M fructose) to a high-quality patch (1M fructose) and decreased even more when larvae moved from a high-quality (1M fructose) to a low-quality (0.1M fructose) patch.

**Figure 4.**
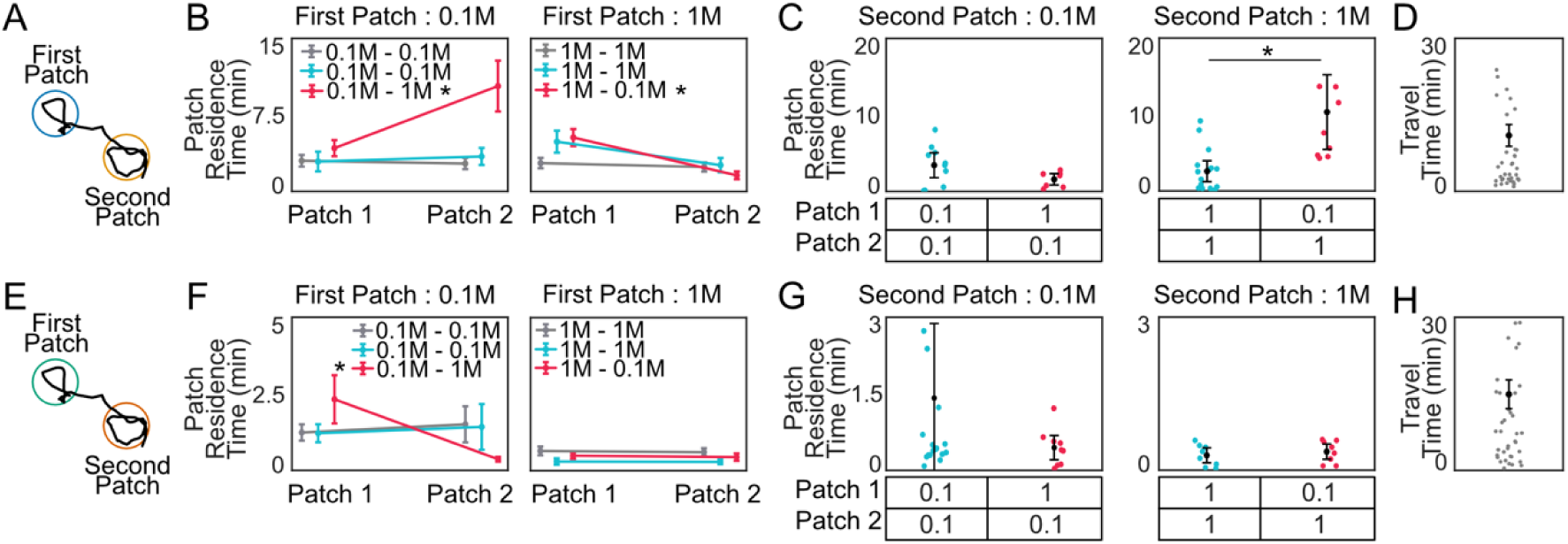
*Drosophila* larvae adapt their behavior based on past foraging experience: (**A**) Schematic illustration of subsequent patch visits by a larva when the patches contained fructose. (**B**) Average patch residence time of larvae on patches 1 and 2 in homogeneous (gray, Fig 1A) or heterogeneous environments (blue for same-type patches, pink for different-type patches; Fig 3A). Data is shown as mean ± SEM (Wilcoxon signed-rank test, * p < 0.05, indicated in the legend). (**C**) Average patch residence time on patch 2 based on the first patch experienced by the larvae in a heterogeneous environment. Dots indicate individuals, and the line represents the mean ± 95% confidence interval. (**D**) Time taken by larvae to travel from patch 1 to patch 2 in a heterogeneous environment. (**E**) Schematic illustration of subsequent patch visits when the patches contained salt. (**F**) Average patch residence time of larvae on patches 1 and 2 in homogeneous (gray, Fig 1A) or heterogeneous environments (blue for same-type patches, pink for different-type patches; Fig 3A). (**G**) Average patch residence time on patch 2 in a heterogeneous environment, based on the first patch experienced by the larvae. Dots indicate individuals; the line represents the mean ± 95% confidence interval (Mann-Whitney U test, * p < 0.05). (**H**) Travel time between patches 1 and 2 in a heterogeneous environment.

To investigate whether past foraging experience influences foraging behavior on the subsequent patch, we quantified the residence time on the second patch based on the first patch encountered by the larvae. We found that there was a reduction in patch residence time on a 0.1M fructose patch when larvae had previously encountered a 1M fructose patch, compared to when they had previously encountered a 0.1M fructose patch (Fig. 4C). Conversely, there was an increase in patch residence time on a 1M fructose patch when larvae had previously encountered a 0.1M fructose patch compared to when they had previously encountered a 1M fructose patch. On average, larvae took approximately 10 minutes to travel from the first to the second patch, indicating that past foraging experience can influence their subsequent foraging behavior over a several-minute timescale (Fig. 4D).

We observed a similar overall pattern in the heterogeneous environment of different salt resource valences compared to the homogeneous environment (Fig. 2). When we compared the patch residence time over subsequent patch visits (Fig. 4E), we found that larvae showed little reduction in patch residence time when transitioning between patches of the same resource valence (Fig. 4F). When larvae moved from an attractive patch (0.1M salt) to an aversive patch (1M salt), there was a decrease in residence time. However, transitioning from an aversive (1M salt) to an attractive (0.1M salt) salt patch did not result in a significant change.

To assess the influence of past foraging experience, we quantified the residence time on the second patch based on the first patch encountered. We found a reduction in residence time on a 0.1M salt patch if larvae had previously encountered a 1M salt patch compared to a 0.1M salt patch (Fig. 4G). However, the residence time on a 1M salt patch was unaffected by prior experience. On average, larvae took approximately 10 minutes to travel between subsequent patches (Fig. 4H).

### A simple integration model recapitulates larval patch foraging behavior

To investigate the dynamics of patch-foraging decisions and test whether larvae integrate evidence during foraging, we used a drift-diffusion model to explain larval behavior in the fructose patch environments. Previous work suggests that integrator models provide a mechanism that implements decision-making during patch foraging under ecologically relevant conditions (12).

Inspired by this approach, we modeled larval behavior in homogeneous environments using a simple drift-diffusion model with μ as the drift variable (Fig. 5A). We found no difference in model fits when the very first patch encounter in the homogeneous environment was included or not (Fig. S4A). To avoid potential effects associated with the initial exposure to a novel patch, we excluded the first encountered patch from the empirical data (Fig. 5B). A model with a single free parameter, drift, recapitulates larval foraging behavior in homogeneous environments (Fig. 5B/C). The addition of a leak term did not improve the model performance (Fig. S4B). Drift values were lower when the patches contained fructose, consistent with the longer patch residence times observed under these conditions (Fig. 5D).

**Figure 5.**
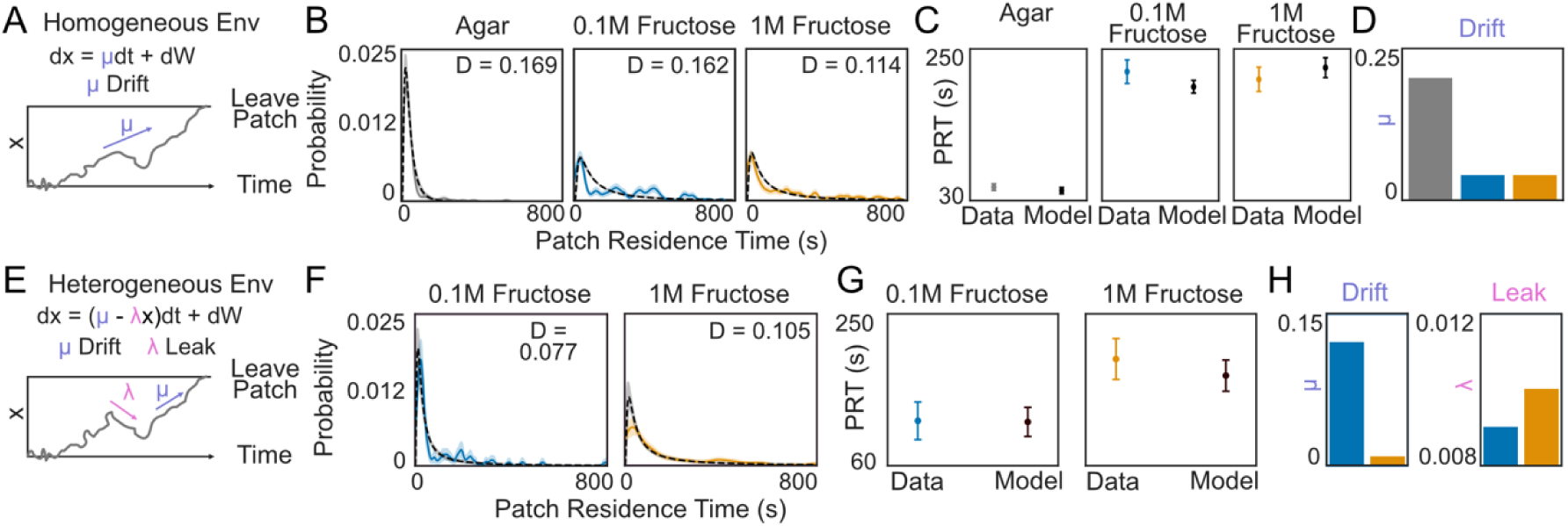
Integration model captures larval patch foraging behavior: (**A**) Implementation of the drift-diffusion model (DDM) for patch leaving in homogeneous environments, with x as the decision variable and μ as the drift variable. (**B**) Model fits for the different substrates (black line: mean fit; shaded area: ± standard deviation; KS test indicates goodness-of-fit). (**C**) Mean ± std of the patch residence times from the empirical data and model prediction. (**D**) Drift rates obtained from the DDM fits (**E**) Implementation of the DDM model for patch leaving in the heterogeneous environment, with x as the decision variable, μ as the drift, and 𝞴 as the leak. (**F**) Model fits for the different substrates (black line: mean fit; shaded area: ± standard deviation; KS test indicates goodness-of-fit). (**G**) Mean ± std of the patch residence times from the empirical data and the model. (**H**) Drift (μ) and leak (λ) obtained from the model fits.

We then extended the drift–diffusion model to describe larval foraging behavior in heterogeneous environments where larvae can encounter two different patch qualities (Fig. 5E). Incorporating a leak term improved the model’s goodness of fit (Fig. 5E, Fig. S4C). Because larvae can only compare resources after encountering both patches, we restricted the analysis to patch-residence data after both patches had been visited by a larva (Fig. 5F), which further improved the model fit (Fig. S4D). The model with drift and leak terms captures larval foraging behavior in heterogeneous environments (Fig. 5G). Drift values were lower, and leak values were higher when the patches contained high-quality fructose, consistent with the longer patch residence times observed for these patches (Fig. 5H).

## Discussion

Our study has provided a systematic analysis of larval foraging behaviors across environments with different structures, offering insights into how *Drosophila* larvae adjust their decision strategies accordingly. Our results demonstrate that in homogeneous patch environments, the foraging behavior of *Drosophila* larvae is shaped by both the quality and valence of a single available resource. In heterogeneous environments with patches of varying quality or valence, larvae flexibly adjust their foraging strategies in response to the resource landscape. Finally, we show that prior experience influences larval foraging decisions and introduce an integration mechanism that tracks the environmental characteristics.

To investigate foraging behaviors, we established a behavioral assay with continuous recording of individual larval behavior over an extended 3-hour timescale, much longer than any previous studies. This extended timescale allows larvae to explore and exploit all available non-depleting resource options. Testing individual larvae allowed for precise identity tracking and eliminated the influence of conspecifics (29).

### Resource quality and valence shape larval foraging

Resource quality influenced the time larvae spent foraging. Larvae spent more time on or in the vicinity of fructose patches at different concentrations (Fig. 1B, Fig. 1C), consistent with their attraction to both 0.1M and 1M fructose (24, 25). The patch residence time did not differ between concentrations, likely because fructose was the sole nutritious resource available in the environment, leading larvae to exploit both concentrations to a similar extent. In contrast, larvae spent significantly less time on agar, indicating that patch exploitation is driven by resource quality rather than the mere presence of a patch border.

Similarly, resource valence also influenced patch residence time. Larvae spent more time on or in the vicinity of patches containing 0.1M salt than on agar or 1M salt concentrations (Fig. 2B, Fig. 2C). In addition, larvae are more likely to return to 0.1M salt patches than to agar and 1M salt patches (Fig. 2D). This is consistent with their attraction to 0.1M salt concentration and perception of 1M salt concentration as punishing (28, 30). Trace amounts of NaCl are essential for various larval physiological functions, such as osmoregulation and neural processes (31, 32). However, ingesting excessive salt negatively affects larval physiology, leading to delayed development and reduced survival (33). This is reflected in the low number of entries made by larvae onto the 1M salt patches (Fig. S2E).

### Resource quality and valence affect larval exploitation and exploration behavior in homogeneous environments

Resource quality modulated larval patch interactions. They had a lower MSD and moved more slowly on fructose and attractive 0.1M salt than on agar or aversive 1M salt (Fig. 1E, S1G, 2E, S2G), consistent with previous findings showing reduced locomotion on sucrose relative to agar (23) and increased larval speed in the presence of aversive stimuli (34). On the patch, chemokinesis can explain the larval behavior of slowing down when sensing an attractive cue, such as fructose or 0.1M salt. However, larvae were also more likely to return to the fructose and 0.1M salt patches than to agar or to 1M salt (Fig. 1D, 2D), consistent with data from a previous study (23). This suggests a sustained interest in nutritive resources and indicates local search behavior, in line with larvae’s tendency to return to and remain near a food stimulus (35). As fructose or salt is not volatile and cannot be detected outside of the patches (Fig. 1F, 2F), larvae must use a different strategy, such as memory or information integration, to successfully return to the patch once they leave it. This is in line with findings in Wosniack et al., who show that a simple model based on chemokinesis cannot explain foraging behavior. Because larvae returned more frequently to attractive patches, indicated by an increase in patch entries across conditions, we calculated patch residence time using a distance threshold that includes the majority of these return trips while excluding time spent outside the patch (Fig. S1D, S2D).

We find no difference in travel time between patches throughout the experiment and across conditions, further indicating that larvae cannot detect fructose or salt outside of the patches (Fig. 1F, S1I, 2F, S2I). A lack of volatile cues might also be the reason larvae leave the attractive food patches more often and only return within a certain radius (3 cm). Testing larval foraging in a food patch environment without olfactory cues allows for better control of stimulus exposure and requires the larvae to rely on and remember recent experiences, suggesting that navigation in such environments relies more on prior foraging experience.

Resource quality and valence influenced larval behavior even when they were off-patch. Larvae had a slightly reduced MSD off-patch when patches contained attractive resources, such as fructose or 0.1M salt (Fig. S1H, 2H). This might be due to local search behavior, as the MSD off-patch included the return trip displacement of larvae outside the patch. The larval MSD off-patch was also reduced when the patches contained 1M salt, likely because salt diffuses over time, creating lower, attractive salt concentration regions near the patch border. Larvae remained closer to high-quality fructose patches and spent less time at the arena borders compared to low-quality fructose patches (Fig. S1J). These results suggest that larvae use information about resource quality to modulate their exploration strategy, biasing their movement to remain close to higher-quality resources. However, larval spatial distribution off-patch did not differ between 0.1M salt and 1M salt patch conditions (Fig. S2J). Salt induces weak attraction in larvae, comparable to low fructose concentrations, likely because larvae require only minimal amounts of salt and can survive under low-salt conditions (36).

### Larvae change their foraging behavior based on available resources

When we presented multiple resources in the arena, larvae were able to differentiate between patch resources and flexibly adjusted their foraging behavior based on the resources available at the patch. Similar to the homogenous assay, larvae stayed longer on the attractive salt patches than on the aversive salt patches (Fig. 3H). However, in the heterogeneous fructose assay, they stayed longer on high-quality fructose patches than on low-quality fructose patches, whereas no difference was observed in the homogeneous environments (Fig. 3D).

Based on these results, we conclude that larvae modulate their foraging behavior by taking prior experience into account. When they encounter patches of the same quality consecutively in a heterogeneous environment, their residence times do not differ, resembling their behavior in a homogeneous environment (Fig. 4B, F). In contrast, after experiencing both resource types, larvae adjust their responses according to resource quality and valence, suggesting that they can compare different resources across patch visits and modify their foraging behavior.

Our data indicate that larvae can maintain a memory of prior resource experience and adapt their foraging behavior at the next patch, even after a travel time of approximately 10 minutes (Fig. 4D, H). This memory is unlikely to be metabolically driven, as residence times remain unchanged when larvae visit subsequent patches of a similar resource quality. The residence time even increases when larvae go from a low-quality fructose patch to a high-quality fructose patch. If metabolic satiety mechanisms played a role, prior exposure to any fructose concentration should always increase fructose satiety rather than enhance subsequent fructose exploitation. Therefore, our findings indicate that larvae perform an experience-dependent comparative evaluation of resource quality.

In changing environments, animals must either learn the absolute values of the available resources (37, 38) or assess their relative quality (39) and continuously update their foraging strategies accordingly. Learning the relative value of resources enables animals to efficiently process information that varies widely in magnitude and has been observed in diverse organisms, including insects (40) and birds (41). *Drosophila* larvae have been shown to perform relative value learning rather than learning the absolute value of rewards or punishments (42). In our experiments, we tested two fructose concentrations; exploring additional concentrations will provide further insight into how larvae evaluate and compare resources.

### A simple integrator model captures larval foraging behavior

While foraging in patchy environments, larvae must balance exploiting the current resource with leaving to search for better alternatives. Theoretical work has suggested that a drift-diffusion model provides a potential neural algorithm through which the decision to leave a patch emerges from the gradual accumulation of information about patch quality over time (12). In homogeneous environments, we could describe larval patch-leaving behavior by a drift–diffusion model with a single free parameter, drift (Fig. 5B/C). The addition of a leak term, which represents the decay or resetting of accumulated information, did not improve the model performance (Fig. S4B), indicating that a simple evidence-accumulation process is sufficient to explain the observed behavior. This suggests that the decision variable does not strongly discount past evidence over time, consistent with the absence of a history effect in patch residence behavior (Fig. 4B). Lower drift values observed in the fructose patches (Fig. 5D) were consistent with longer residence times, suggesting that nutritious resources slow the accumulation of patch-leaving evidence, thereby prolonging exploitation.

In heterogeneous environments, however, the introduction of a leak term improved model accuracy for describing larval behavior (Fig. 5F/G). This improvement in model fit implies that larvae integrate present sensory information with a decaying influence based on prior patch experiences, captured here by the leak term. The observed pattern of lower drift and higher leak in high-quality fructose conditions (Fig. 5H) suggests slower accumulation of sensory evidence about patch quality, together with faster decay of previously integrated information. This dynamic tuning of drift and leak, which depends on foraging history, may enable larvae to balance exploitation of currently favorable resources with exploration of novel patches within the environment. Together, our findings suggest that larval foraging decisions in changing environments can emerge from a simple integration process whose parameters are modulated by environmental statistics and sensory experience.

Establishing this foraging paradigm in *Drosophila* larvae will allow us to leverage powerful genetic tools together with the whole-brain connectome to decode the neural circuits underlying foraging decisions. Larvae frequently return to the same patch, and such persistent behavior to food has been linked to octopaminergic neuromodulation in *C. elegans* and adult *Drosophila* (43, 44). We find that larvae can compare resources and shape their foraging decisions based on prior experience. In adult *Drosophila*, dopamine neurons have been shown to encode the relative values of attractive and aversive stimuli (45, 46). Comparable dopaminergic circuits are present in larvae (47, 48), suggesting that similar mechanisms may support relative value coding during larval foraging. Our modeling results further provide a quantitative framework for identifying the parameters governing evidence integration during patch exploitation, raising the question of how such computations are implemented in neural circuits. Evidence integration has been proposed to involve mushroom body circuits in adults, where neural activity can reflect accumulated sensory information during decision-making (49, 50). Together, combining this behavioral paradigm with circuit-level perturbations will allow us to directly test how larvae compare resource values and implement the integrative computations predicted by our model.

## Materials and Methods

### Animal stocks and husbandry

Canton-S flies were reared on a standard cornmeal diet and maintained in incubators at 25 °C and 60% relative humidity under a 12 h light/12 h dark cycle. Adult flies were allowed to oviposit for 48 h, after which they were removed from the vials. The eggs were allowed to develop for 4–6 days. Larvae at the early third-instar stage were used for experiments.

### Behavioral experiments

The experiments were conducted in a 25 cm x 25 cm assay arena filled with 100ml of a 2% agarose substrate. After the agar solidified, four circular holes of radius 1.75cm were made using Falcon tubes, positioned at equal distances of 6.25 cm from the center of the arena. To make patches, the agar within these holes was removed and replaced with agar, fructose, or salt solutions filled to the same height as the surrounding agar substrate.

We added 1.8g or 18g of fructose to 100ml of a 2% agarose solution to prepare 0.1M and 1M fructose solutions, respectively. To make 0.1M and 1M salt solutions, we added 0.58g or 5.8g of salt to 100ml of a 2% agarose solution, respectively. All four patches were filled with the same substrate to make homogeneous arenas. To make heterogeneous arenas, two neighbouring patches were filled with 0.1M solutions, while the remaining two patches were filled with 1M solutions. All arenas were used for behavioral experiments immediately after preparation.

All arenas were maintained at a constant temperature of 25°C and 60% humidity and placed in a light-tight box. For all behavioral experiments, early third-instar larvae of a similar size were used (4-6 days after egg laying). The larvae were removed from the fly vials and washed with distilled water to remove all traces of food.

A single larva was placed in the middle of the assay arena, and its behavior was recorded for 3 hours with a Basler camera (acA2040-90umNIR) and lens (Kowa Lens LM16HC F1.4 f15mm1”) at 1fps with a red light filter (Edmund optics #89-837) positioned above the arena.

### Behavioral Analysis

Videos were analyzed using the freely available tracking software TRex (51) to obtain larval position (X/Y) over the 3-hour experiment.

### Patch residence time

To quantify patch residence time, we calculated the duration of each visit to a patch. Upon leaving the patch, we characterized post-exit behavior by measuring the maximum displacement of the larva from the patch edge and the return time for every trip. Given the correlation between return time and maximum displacement, we continued with displacement as a measure of post-exit behavior. Because the majority of larvae returned to the patch after trips limited to ≤ 3 cm from the patch edge, we defined this as the threshold for calculating patch residence time. While calculating patch residence time, a patch visit was defined as a continuous period when the larva was either on the patch or made return trips of less than 3 cm from the patch’s edge. However, only the time spent on the patch was included in the patch residence time calculation.

To analyse patch residence times across subsequent patch visits, we focused on the first two patch transitions between patches of different quality to increase the sample size. This included the transition from the first to the second patch and, in cases where the patch type remained the same, from the second to the third patch.

### Larval movement analysis

To quantify larval dispersal, we calculated the mean squared displacement (MSD) for specific time lags (*τ*) as follows:

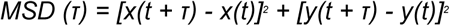

At each frame, the MSD was calculated for all past *τ* frames, if possible. For each *τ*, the average MSD was calculated as the mean of all MSD values for that *τ*. We then computed the MSDs separately for on-patch and off-patch behaviors by averaging the values across all larvae for each condition. Additionally, we quantified larval speed both on and off patches. Finally, we quantified travel time as the duration between patch exit and arrival at the same or subsequent patch following the first patch exit using the displacement threshold defined above.

### Drift-diffusion model

A drift-diffusion model was implemented using an Ornstein-Uhlenbeck process to simulate the foraging dynamics in homogeneous environments as follows:

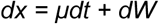

where x is the decision variable, *μ* is the drift parameter, and W denotes a Wiener process, representing stochastic noise.

To model the foraging dynamics in heterogeneous environments, a leak (*λ*) was incorporated into the above model as follows:

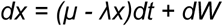

For each empirical dataset, bootstrapping was performed by resampling patch residence times 1000 times to estimate the variability of the model parameters.

### Statistical analysis

To determine significant differences between the experimental groups, we performed a non-parametric Mann-Whitney U-test (Mann-Whitney U test = * p < 0.05). The test was conducted using the substrate as the grouping factor. To compare patch residence times over subsequent patches, we performed a non-parametric Wilcoxon signed-rank test for paired data. To evaluate the goodness of fit of the model, we used a non-parametric Kolmogorov–Smirnov (KS) goodness-of-fit and reported the D statistic, where lower D values indicate a better fit (Dataset S1).

## Author Contributions

Conceptualization: A.M., A.E.H., K.V. Methodology: A.M. Software: A.M. Validation: A.M. Formal analysis: A.M. Investigation: A.M., K.T. Data curation: A.M. Resources: K.V. Writing - original draft: A.M., A.E.H., K.V. Writing - review and editing: A.M., A.E.H., K.V. Visualization: A.M. Supervision: A.E.H., K.V. Project administration: K.V. Funding acquisition: K.V.

## Acknowledgments

We would like to thank Andreas Thum and Iain Couzin for valuable discussions. A.M., A.E.H., and K.V. were supported by the DFG German Research Foundation (EXC 2117-422037984). A.E.H. was also supported by the Human Frontiers Science Foundation Grant (RGP006/2025). K.T. was supported by the Erasmus+ programme. Open access funding provided by Max Planck Society.

## Supplementary figures

**Fig. S1.**
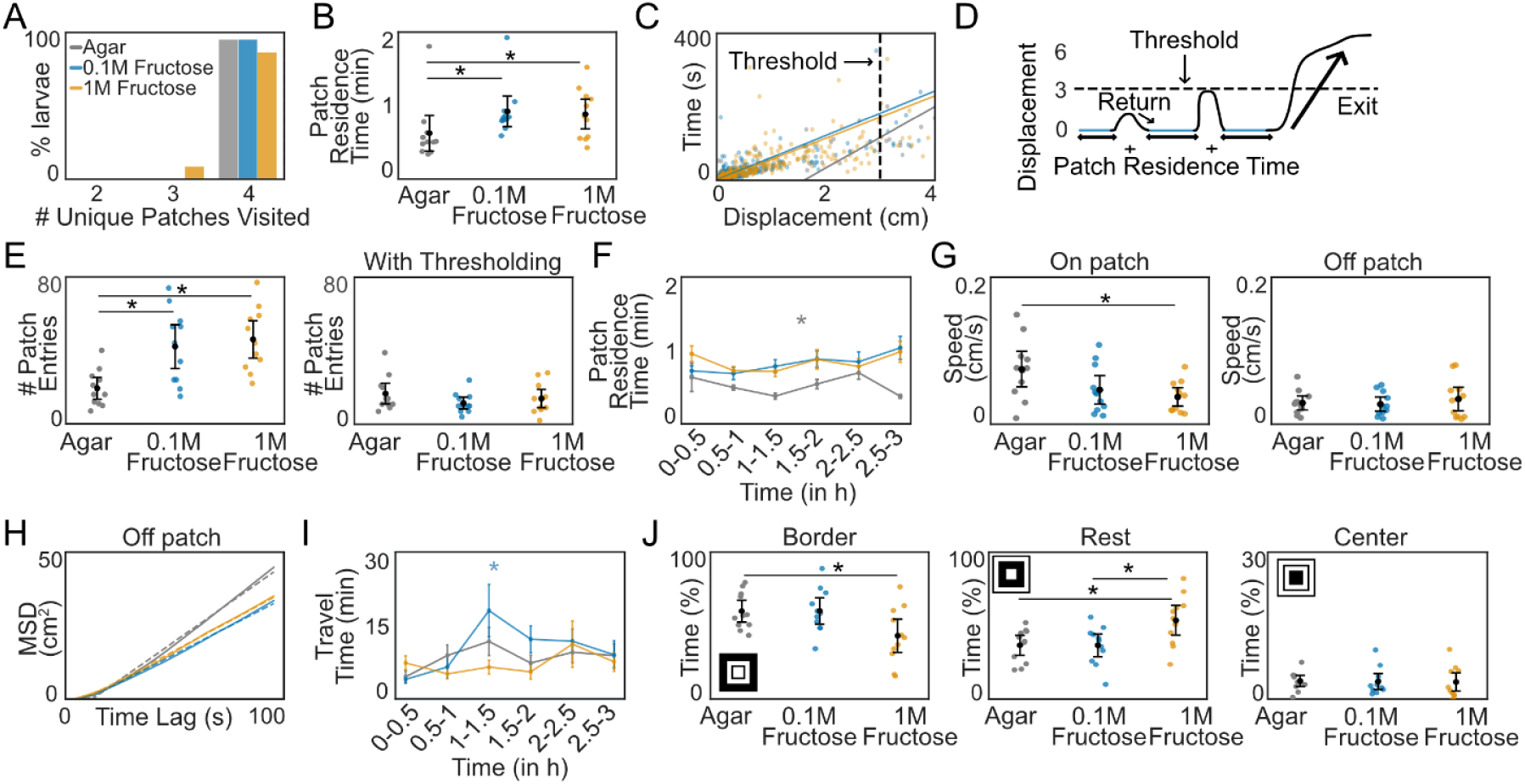
Analysis of larval behavior in homogeneous environments of different resource qualities: (A) Number of unique patches visited by larvae when the patches contained agar (gray), 0.1M fructose (blue), and 1M fructose (yellow). (B) Average patch residence time of larvae. Dots indicate individual larvae; the line represents the mean ± 95% confidence interval (Mann-Whitney U test = * p < 0.05). (C) Correlation between maximum displacement from the patch edge and the return time to the patch. Each dot represents an individual trip, and the line indicates the linear regression fit. (D) Schematic illustrating the calculation of patch residence time after applying a distance threshold. (E) Number of patch entries by each larva before and after thresholding. (F) Thresholded average patch residence time in 30 min interval (Kruskal-Wallis test = * p < 0.05). (G) Speed of larvae on- and off-patch. (H) MSD of larvae when they were off-patch. The dotted line indicates the linear fit to the MSD. (I) Travel time in 30 min intervals (Kruskal-Wallis test = * p <0.05). (J) Time spent by larvae when they were off-patch at the border, in regions near the patches and in the center of the arena.

**Fig. S2.**
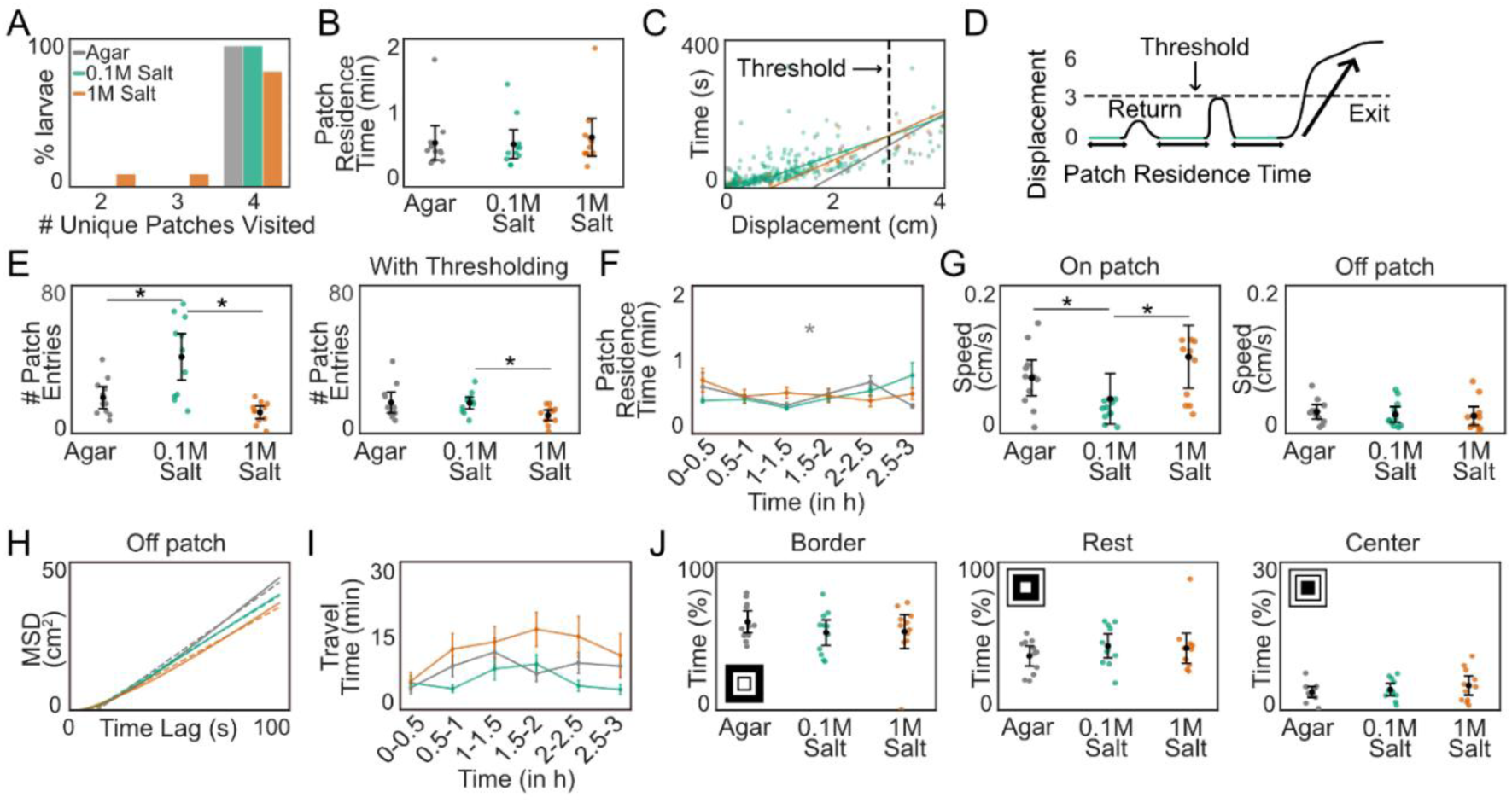
Analysis of larval behavior in homogeneous environments of different resource valences: (A) Number of unique patches visited by larvae when the patches contained agar (gray), 0.1M salt (green) and 1M salt (orange). (B) Average patch residence time of larvae. Dots indicate individual larvae; the line represents the mean ± 95% confidence interval (Mann-Whitney U test = * p < 0.05). (C) Correlation between maximum displacement from the patch edge and the return time to the patch. Each dot represents an individual trip and the line indicates the linear regression fit. (D) Schematic illustrating the calculation of patch residence time after applying a distance threshold. (E) Number of patch entries by a larva before and after thresholding. (F) Thresholded average patch residence time over 30-min intervals (Kruskal-Wallis test = * p < 0.05). (G) Larval speed on- and off-patch (H) MSD of larvae when they are off-patch; the dotted line indicates the linear fit to the MSD. (I) Travel time between patches at 30-min intervals (Kruskal-Wallis test). (J) Time spent by larvae when they were off-patch at the border, in regions near the patches and in the center of the arena.

**Fig. S3.**
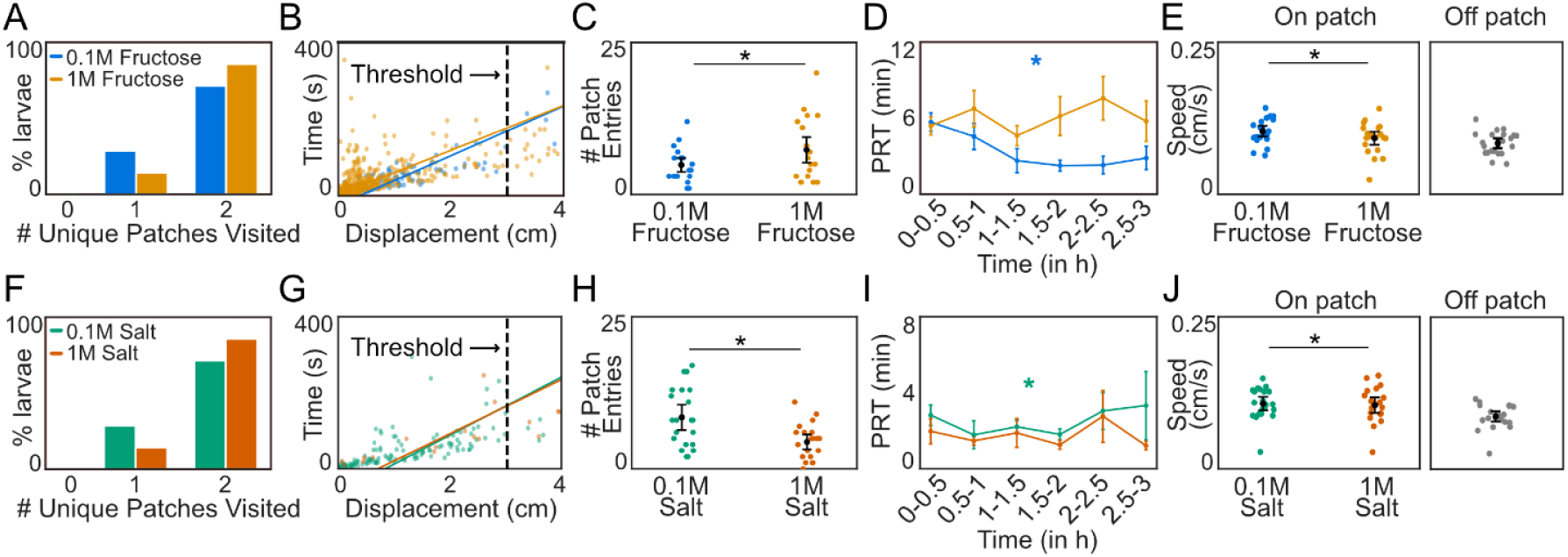
Analysis of larval behavior in a heterogeneous environment of different resource qualities and valences: (**A**) Number of unique patches visited by larvae when two patches contained 0.1M fructose (light blue) and 1M fructose (blue). (**B**) Correlation between maximum displacement from the patch edge and the return time to the patch. Each dot represents an individual trip, and the line indicates the linear regression fit. (**C**) Number of patch entries by a larva after thresholding. Dots indicate individual larvae; the line represents the mean ± 95% confidence interval (Mann-Whitney U test = * p < 0.05). (**D**) Thresholded patch residence time over time in 30 min intervals (Kruskal-Wallis test = * p < 0.05). (**E**) Speed of larvae on- and off-patch. (**F**) Number of unique patches visited by larvae when two patches contained 0.1M salt (light red) and 1M salt (red). (**G**) Correlation between maximum displacement from the patch edge and the return time to the patch. (**H**) Number of patch entries by larvae after thresholding. (**I**) Thresholded patch residence time over time in 30 min intervals (Kruskal-Wallis test = * p < 0.05). (**J**) Larval speed on- and off-patch.

**Fig. S4.**
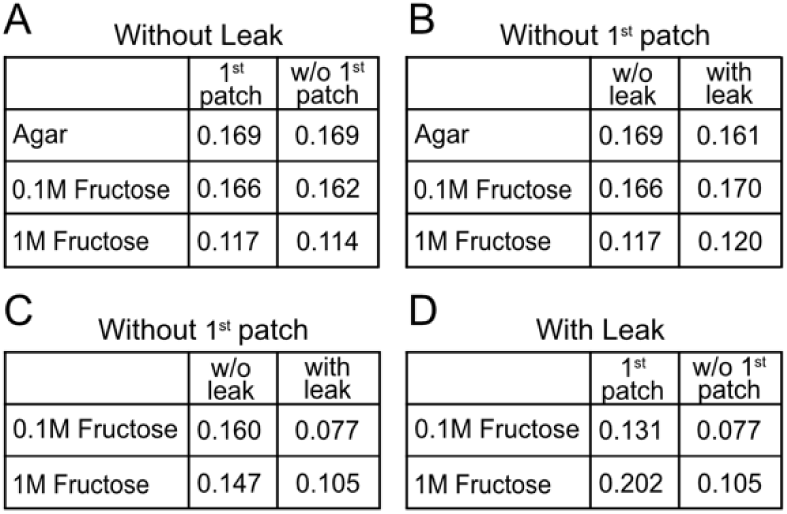
Analysis of DDM model parameters: (**A**) Goodness-of-fit values for the drift-diffusion model for patch-leaving decisions, with and without inclusion of the first encountered patch, with x as the decision variable and μ as the drift, based on the empirical data from Fig. 5A. (**B**) Goodness-of-fit values for the drift-diffusion model for patch-leaving decisions excluding the first encountered patch, evaluated with and without λ as the leak, with x as the decision variable and μ as the drift, based on the empirical data from Fig. 5A.(**C**) Goodness-of-fit values for the drift-diffusion model for patch-leaving decisions excluding encounters prior to the exploration of both patch types, evaluated with and without λ as the leak, with x as the decision variable and μ as the drift, applied to the empirical data from Fig. 5E. (**D**) Goodness-of-fit values for the drift-diffusion model for patch-leaving decisions, with and without inclusion of the first encountered patch of both types, with x as the decision variable, μ as the drift and λ as the leak, applied to the empirical data from Fig. 5E.

## Supplementary Videos

**Movie S1.** Foraging behavior of an individual larva exploring an arena containing four **0.1M fructose** patches (in blue) for 3 hours (shown at 10x).

**Movie S2.** Foraging behavior of an individual larva exploring an arena containing four **1M fructose** patches (in yellow) for 3 hours (shown at 10x).

**Movie S3.** Foraging behavior of an individual larva exploring an arena containing four **agar** patches (in gray) for 3 hours (shown at 10x).

**Movie S4.** Foraging behavior of an individual larva exploring an arena containing four **0.1M salt** patches (in green) for 3 hours (shown at 10x).

**Movie S5.** Foraging behavior of an individual larva exploring an arena containing four **1M salt** patches (in orange) for 3 hours (shown at 10x).

**Movie S6.** Foraging behavior of an individual larva exploring an arena containing two **0.1M fructose** patches (in blue) and two **1M fructose** patches (in yellow) for 3 hours (shown at 10x).

**Movie S7.** Foraging behavior of an individual larva exploring an arena containing two **0.1M salt** patches (in green) and two **1M salt** patches (in orange) for 3 hours (shown at 10x).

